# A coordinated function of lncRNA HOTTIP and miRNA-196b underpinning leukemogenesis by targeting Fas signaling

**DOI:** 10.1101/2021.07.15.452559

**Authors:** Ajeet P. Singh, Huacheng Luo, Meghana Matur, Melanie Eshelman, Arati Sharma, Suming Huang

## Abstract

MicroRNAs (miRNAs) may modulate more than 60% of human coding genes and act as negative regulators, while long non-coding RNAs (lncRNAs) regulate gene expression on multiple levels by interacting with chromatin, functional proteins, and RNAs such as mRNAs and microRNAs. However, the crosstalk between lncRNA *HOTTIP* and miRNAs in leukemogenesis remains elusive. Using combined integrated analyses of global miRNA expression profiling and state-of-the-art genomic analyses of chromatin such as ChIRP-seq., (genome wide *HOTTIP* binding analysis), ChIP-seq., and ATAC-seq., we found that miRNA genes are directly controlled by *HOTTIP.* Specifically, the HOX cluster miRNAs (miR-196a, miR-196b, miR-10a and miR-10b), located *cis* & *trans*, were most dramatically regulated and significantly decreased in *HOTTIP*^-/-^ AML cells. *HOTTIP* bound to the miR-196b promoter, and *HOTTIP* deletion reduced chromatin accessibility and enrichment of active histone modifications at HOX cluster associated miRNAs in AML cells, while reactivation of *HOTTIP* restored miR gene expression and chromatin accessibility in the CTCF-boundary-attenuated AML cells. Inactivation of *HOTTIP* or miR-196b promotes apoptosis by altering the chromatin signature at the *FAS* promoter and increasing *FAS* expression. Transplantation of miR-196b knockdown MOLM13 cells in NSG mice increased overall survival compared to wild-type cells. Thus, *HOTTIP* remodels the chromatin architecture around miRNAs to promote their transcription, consequently repressing tumor suppressors and promoting leukemogenesis.

## Introduction

Non-coding RNAs are emerging as important regulators of gene expression in multiple cellular processes, especially in cancer^1^. In particular, long non-coding RNAs (lncRNAs) are involved in regulating chromatin dynamics, gene expression, cell growth, differentiation, and development^2^. Overexpression of *HOXA9* is a dominant factor driving certain subtypes of human acute myeloid leukemia (AML)^3^. Moreover, the aberrant activation of posterior HOXA gene, *HOXA9,* and its cofactor, *MEIS1*, following a variety of genetic alterations, including MLL-translocations, NUP98-fusion, CDX dysregulation, and monocytic leukemia zinc-finger fusion, was associated with poor prognosis and treatment response^4^. LncRNAs regulate transcription through recruiting histone modifiers and chromatin remodeling factors that play active roles in various aspects of development and disease state^5–10^. The lncRNA *HOTTIP*, located at the posterior end of HOXA gene cluster, acts as a scaffold to recruit the WDR5-MLL-complex to the promoters of posterior HOXA genes and positively regulates their expression in normal hematopoiesis and AML leukemogenesis^5,11^. In contrast, loss of *HOTTIP* strongly inhibits posterior (*HOXA9*-*HOXA13*) compared to anterior HOXA gene expression (e.g., *HOXA1-HOXA7*)^11^.

*HOTTIP* provides a basis for transcriptional activation and three-dimensional (3D) chromatin organization in the 5’ HOXA gene loci by acting downstream of the CBS7/9 boundary (CTCF binding site located between *HOXA7* and *HOXA9*)^11^. Our previous data indicated that depleting *HOTTIP* reduces active histone marks (H3K4me3 and H3K79me2) and enhances repressive histone marks (H3K27me3) resulting in a switch from an active to a repressive chromatin state in the promoter region of the 5’ HOXA genes in MOLM13 cells. Genome-wide binding analysis of *HOTTIP* lncRNA using chromatin isolation by RNA purification combining deep sequencing (ChIRP-seq) revealed that *HOTTIP* directly regulates its target genes through *cis* and *trans* binding at *HOXA9-13*, *RUNX1* and *MEIS1* genes. Consequently, *HOTTIP* also binds in *cis* and *trans* regulatory patterns to non-coding regions and certain annotated genes involved in chromatin organization, hematopoiesis, myeloid cell differentiation, cell-cycle progression, and JAK-STAT and WNT signaling pathways. These findings suggested that *HOTTIP* lncRNA might control gene expression by interacting with and regulating non-coding regulatory elements such as microRNAs (miRNAs) in *cis* and *trans*^11^.

Growing evidence indicates that non-coding RNAs, in particular lncRNAs and microRNAs, regulate one another and cooperate to influence the levels of target mRNAs in a cell-type specific manner^12^. LncRNAs process, interact with, and regulate miRNAs at both transcriptional and post-transcriptional levels^13^. LncRNAs can function as miRNA sponges, acting as decoys to impair the functional interaction of a miRNA and its target mRNA, thereby preventing suppression of gene expression^14^. Additionally, lncRNAs can be precursors of miRNAs and regulate miRNA biogenesis at different points. Primary miRNAs (pri-miRNAs) are transcribed in the genome (i) either within the body of another gene and often their expression is linked to the expression of the parent transcript, or (ii) from independent miRNA genes, similar to mRNA, where transcription is primarily controlled by RNA polymerase II-driven promoters^15^. However, the molecular mechanisms, particularly those mediated by lncRNAs, regulating miRNAs transcription remains elusive.

MicroRNAs (miRNAs) are 20-22 nucleotide non-coding RNAs that inhibit target gene expression post-transcriptionally either by translational repression or mRNA degradation via binding at complementary seed sequences mainly located at the 3’ UTR. Widespread dysregulation of certain miRNAs associated with hematological malignancies, including acute myeloid leukemia^16^. Besides the role of lncRNA *HOTTIP* in regulating hematopoietic genes, miRNAs also control leukemic and tumor suppressor gene expression^17^. In particular, miR-196b, which is located adjacent to *HOXA9*, targets *HOXA9* and its cofactor *MEIS1^18^*. It also targets proapoptotic factor *FAS*^19^, suggesting a double-edged sword (miR-196b) that could simultaneously repress both the oncogenic and tumor suppressor target genes. Recent studies have shown the importance of miR-196b in AML; however, the mechanism of its transcriptional regulation remains unknown in AML. Thus, in this study we characterize the mechanisms by which *HOTTIP* controls the expression and function of *HOXA9* and *FAS* through miR-196b in human AML.

## Results

### *HOTTIP* lncRNA differentially regulates microRNAs in AML

To define the mechanism by which *HOTTIP* regulates gene expression in AML, we compared genome wide microRNA expression patterns between wild-type (WT) and *HOTTIP*^*-/-*^ MOLM13 AML cells by performing small-RNA-sequencing (smRNA-seq) analysis (Figure 1A). Differential expression analysis shows that a total 569 miRNAs changed more than 2-fold in *HOTTIP*^*-/-*^ cells compared to WT cells, including 333 down-regulated miRNAs and 236 upregulated miRNAs (Figure 1B, C). These altered miRNAs in *HOTTIP*^-/-^ cells play vital roles in molecular and cellular processes, including hematopoiesis and leukemogenesis, suggesting that *HOTTIP* controls AML pathogenesis by regulating miRNAs. Gene ontology (GO) analysis revealed that *HOTTIP-*regulated miRNAs were involved in many biological processes including the metabolic process, regulation of gene expression, transcription, and RNA processing (Figure 1D).

**Figure 1.**
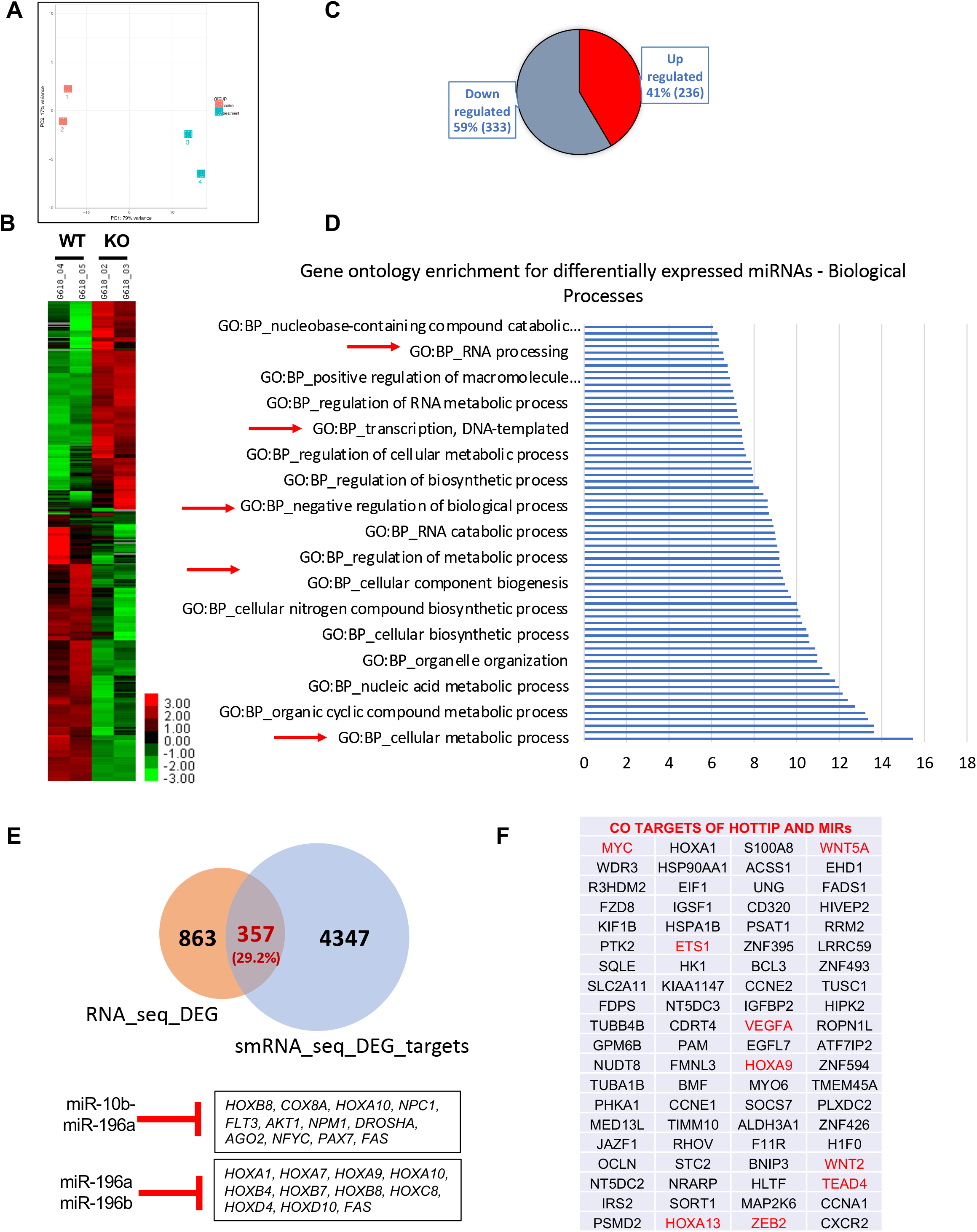
LncRNA-*HOTTIP* regulates miRNAs target genes control leukemogenic program in AML cells. (A) Principal component analysis (PCA) of the miRNAs differentially expressed in *HOTTIP*^-/-^ vs WT AML cells. (B) Heatmap of miRNAs changed >2-fold up- and down upon *HOTTIP*^-/-^ by small-RNA sequencing (smRNA-seq). (C) Select number of mRNAs up- and down regulated in *HOTTIP*^-/-^ MOLM13 cells. (D) GO (Gene Ontology) analysis of miRNAs differentially expressed in *HOTTIP*^-/-^ cells. (E) Number of genes regulated by either miRNAs or *HOTTIP*. (F) Select number of DEGs in *HOTTIP*^-/-^ are the direct target of miRNAs that are regulated by *HOTTIP*.

Next, we sought to determine whether *HOTTIP-*regulated miRNAs control the expression of *HOTTIP-*regulated mRNAs. *In silico* analysis revealed that *HOTTIP*-regulated miRNAs are predicted to directly target 4347 mRNAs. We compared these with 863 mRNAs that we previously demonstrated are transcriptionally regulated by *HOTTPI^11^.* Strikingly, 357 genes are co-regulated by both *HOTTIP*-associated miRNAs and *HOTTIP* itself in AML, including hematopoietic and leukemic signature genes like HOXA factors (e.g., *HOXA9*, *HOXA13*), *MYC*, *ETS1*, *VEGFA*, *ZEB2*, *TEAD4*, *WNT5a*, *WNT2* and *FAS* (Figure 1E, F).

### Depletion of*HOTTIP* inhibits the HOX clusters’ miRNAs in *cis – trans*

Within the HOX cluster several non-coding transcripts, including miRNAs, are transcribed. Six miRNAs are transcribed within the intergenic regions of the HOX clusters (Figure 2A). Loss of *HOTTIP* impaired posterior HOXA gene expression (e. g. *HOX13-HOXA9*) but did not affect anterior *HOXA1-7* genes, HOXB, HOXC, or HOXD genes^11^. We asked whether depletion of *HOTTIP* affects miRNAs that are transcribed in either *cis* or *trans* in the intergenic region of the HOX cluster genes. Disruption of *HOTTIP* inhibits several HOX cluster-related miRNAs, including miR-196b-5p, miR-196a-5p, miR-10b-5p, and miR-10a-5p/3p (Figure 2B), suggesting these miRNAs are positively regulated by *HOTTIP* in AML. Interestingly, while HOX cluster-associated miRNAs were altered, several HOX genes targeted by these miRNAs remained unaltered in *HOTTIP*^-/-^ cells (Figure 1E). We performed correlation analysis of *HOXA9*, its cofactor *MEIS1*, miR-196b and *HOTTIP* according to the RNA-seq dataset of acute myeloid leukemia samples^18^. Our data analysis indicated that miR-196b correlates with expression of *HOTTIP*, posterior *HOXA* genes, *MEIS1,* and *PBX3*, which are aberrantly overexpressed in MLL-associated AML (Figure 2C, D). In contrast, pro-apoptotic genes, including *FAS*, *CASP3,* and *CASP9,* were downregulated in MOLM13 cells relative to miR-196b (Figure 2D). Additionally, miR-196a/b was enriched in *MLL-AF9* MOLM13 cells compared to non-MLL-rearranged leukemia cells such as KASUMI-1/t (8;21) (Figure 2E). However, the question remains whether miR-196b targets pro leukemic gene *HOXA9* in non-MLL-rearranged leukemia cells. The co-expression of *HOTTIP* and miR-196b and their inverse co-relation with pro-apoptotic genes suggests that *HOTTIP* and miR-196b may co-target these genes in *MLL-AF9* rearranged MOLM13 AML cells. Likewise, perhaps *HOTTIP* counteracts miR196b mediated repression of *HOXA9* in AML.

**Figure 2.**
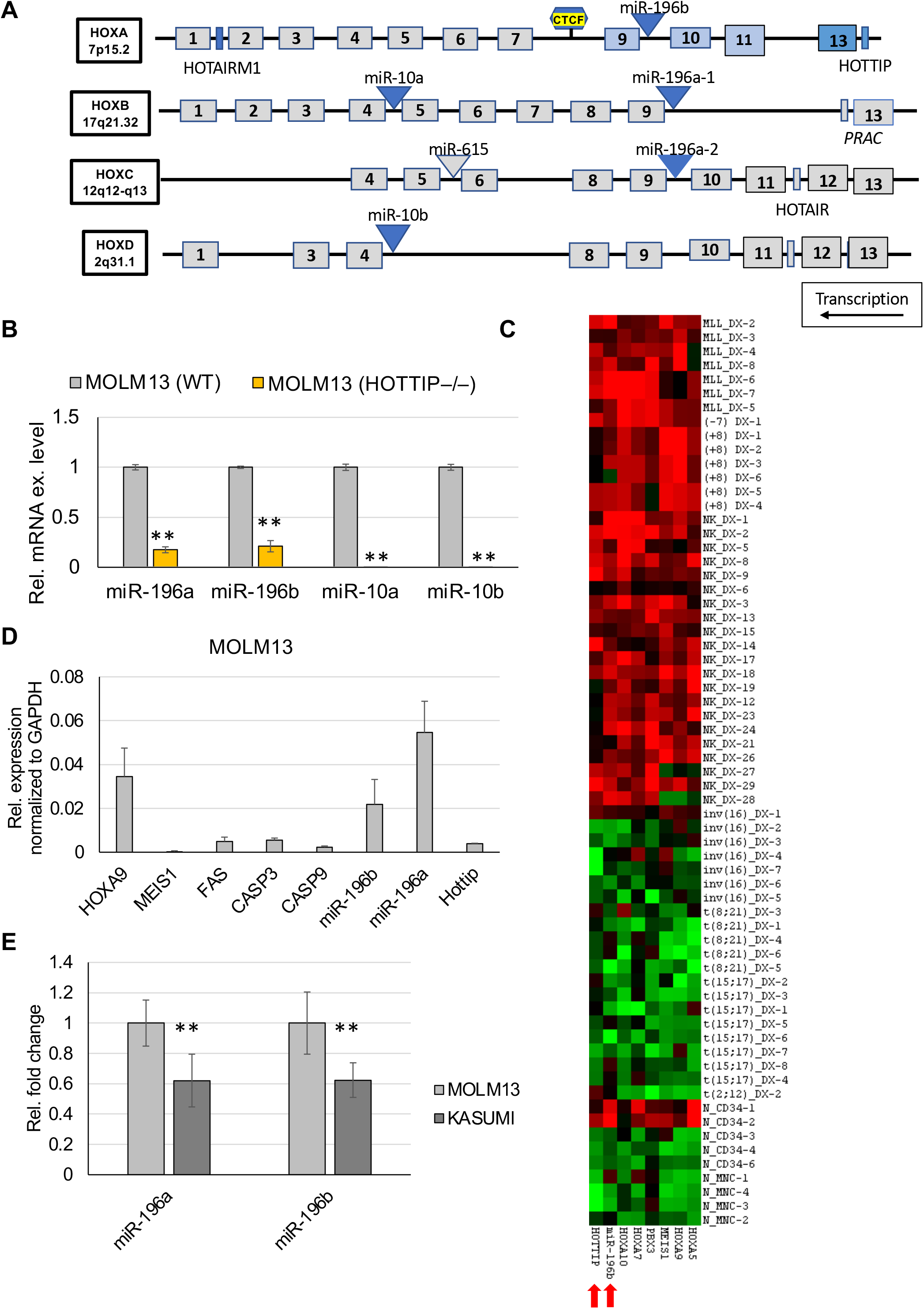
*HOTTIP*^-/-^ perturbs HOX-cluster miRNAs, mediate oncogenic program. (A) Schematics represent HOX cluster coding and non-coding genes. HOX genes are indicated as numbered boxes, miRNAs are shown as triangles and lncRNAs are presented as rectangles. (B,) Expression level of HOX cluster miRNAs in WT and *HOTTIP*^-/-^ MOLM13 cells. (C) Expression correlation between miR-196b, *HOTTIP*, and *HOXA* genes in *de novo* AML and normal control dataset. (D) Relative expression level of the indicated mRNAs, miRNAs and lncRNA in the MOLM13 cells. (F) Relative expression level of the indicated HOX cluster miRNAs in MOLM13 and KASUMI cells.

### *HOTTIP* established chromatin signature on HOX clusters related miRNAs loci

To further define the mechanism by which *HOTTIP* regulates miRNA expression, we employed ChIRP-sequencing (chromatin isolation by RNA purification) to identify *HOTTIP* binding genome-wide^11^. Our previous *HOTTIP* CHIRP-seq data indicated that *HOTTIP* mainly binds to non-coding regions, including promoter and intergenic regions. The question then arises whether *HOTTIP* binds to regulatory loci of miRNA to induce the differential expression we observed in *HOTTIP*^-/-^ compared to WT MOLM13 cells. Our data suggests that loss of *HOTTIP* greatly reduces its binding at the upstream promoter region of miR-196b compared to WT MOLM13 cells (Figure 3A), which is correlated with attenuated expression of miR-196b upon *HOTTIP*-depletion (Figure 2C). Furthermore, *de novo* binding motif analysis revealed that the *HOTTIP* bound genomic locus at miR-196b is occupied by transcription factors, CTCF, GATA-2, *MYC* and *CEBPB*, as well as many other chromatin remodeling factors (Supplementary Figure 1).

**Figure 3.**
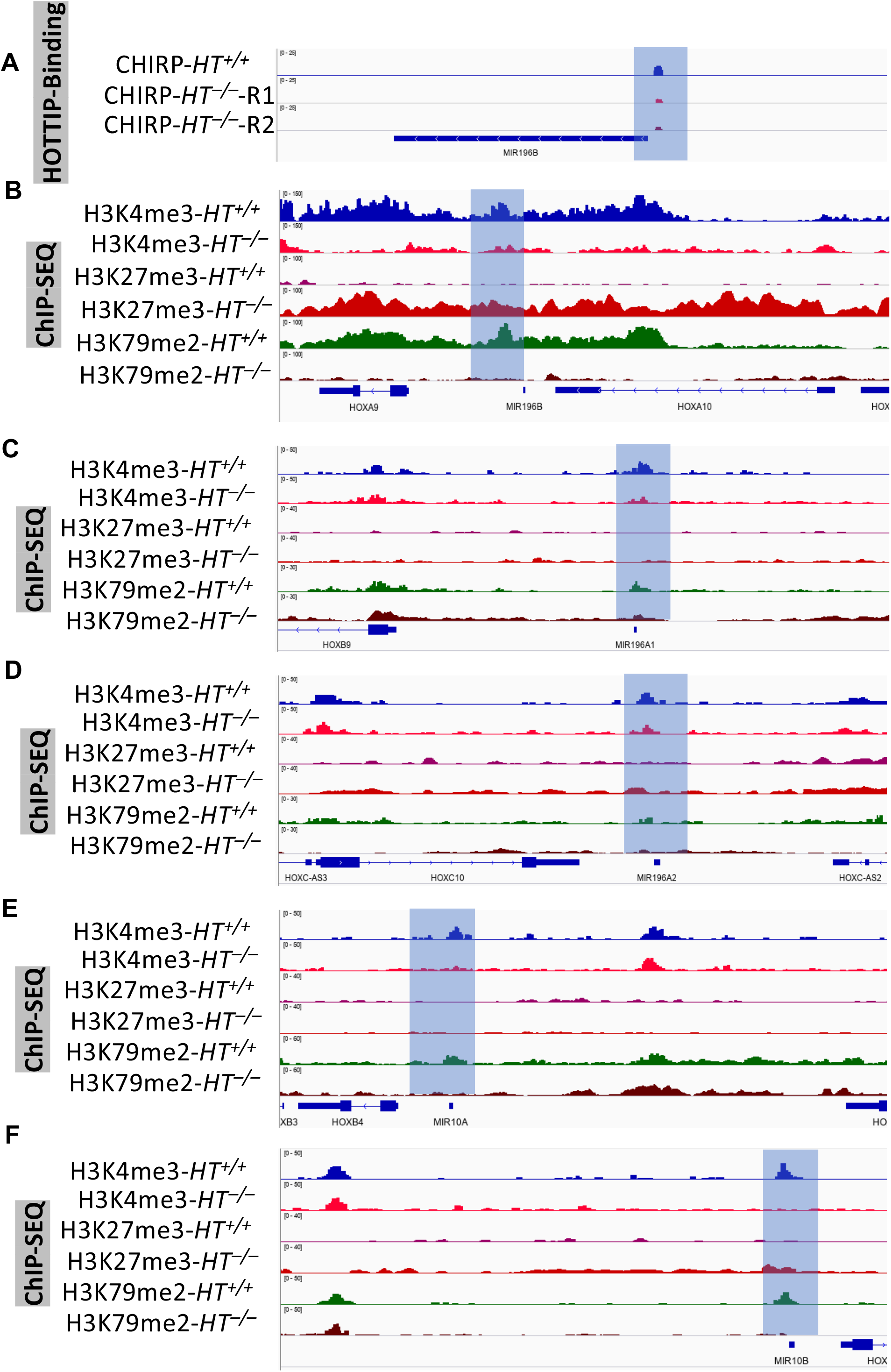
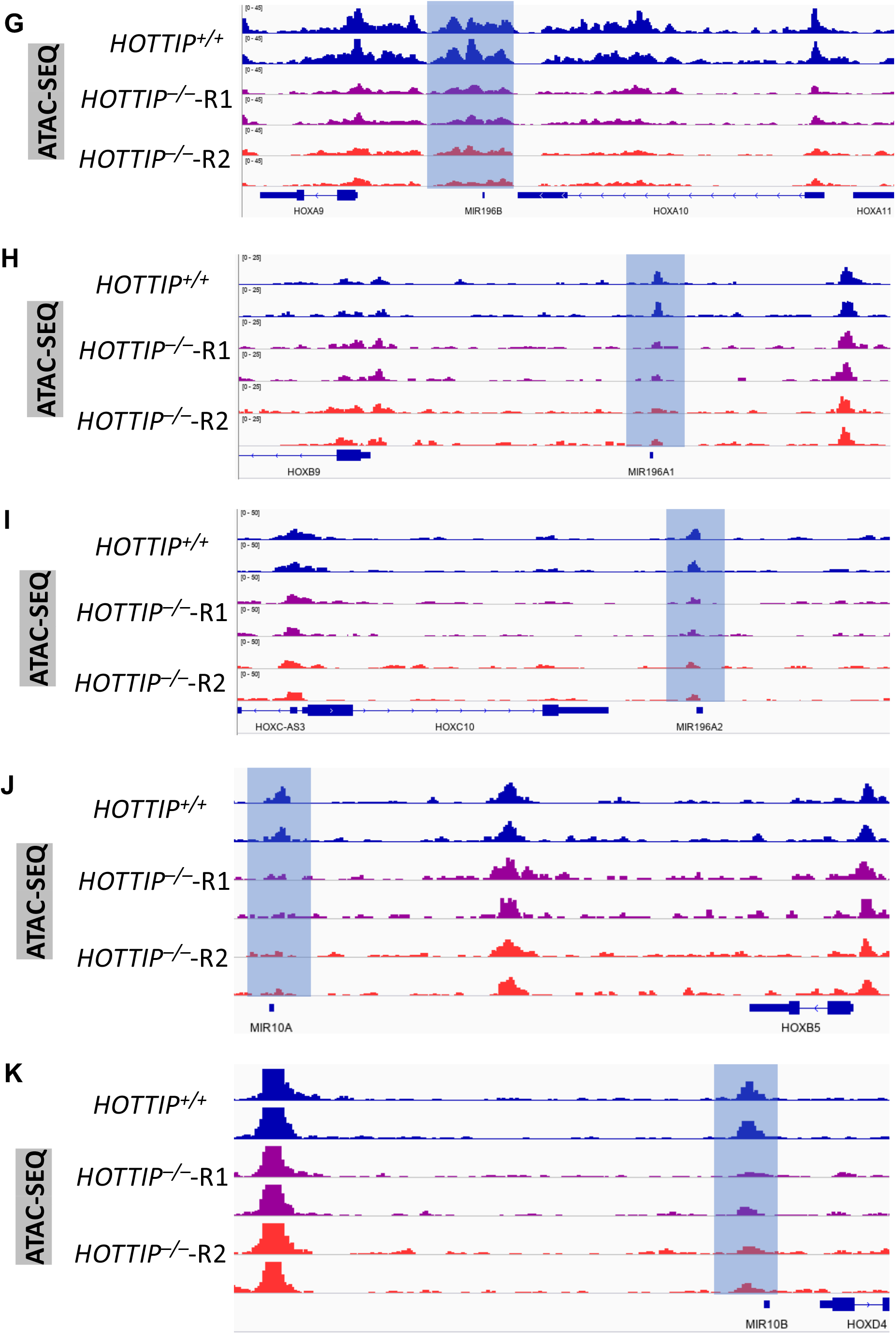
*HOTTIP* controls the epigenetic modifications of HOX cluster miRNAs. (A) ChIRP-seq analysis shows *HOTTIP* binding on promoter region of miR196b in WT and *HOTTIP*^-/-^ MOLM13 cells. (B, C, D, E, F) Enrichment of the active histone marks (H3K4me3 and H3K79me2) and repressive histone mark (H3K27me3) on genomic loci of the HOX cluster miRNAs. (G, H, I, J, K) ATAC-seq analysis shows chromatin accessibility on the HOX-cluster miRNAs upon *HOTTIP*^-/-^ in MOLM13 cells, and WT cells.

Next, we asked whether *HOTTIP* controls the chromatin signature on its targeted miRNA loci, using ChIP-seq (chromatin immunoprecipitation sequencing) for histone modifications (H3K4me3, H3K79me2; active; and H3K27me3; repressive) and ATAC-seq (assay for transposase-accessible chromatin using sequencing) to compare chromatin signatures between WT and *HOTTIP*^-/-^ MOLM13 cells^11^. Depletion of *HOTTIP* resulted in marked decreases in H3K4me3 and H3K79me2 enrichment, while H3K27me3 levels were elevated on the genomic loci of all four HOX cluster miRNAs (Figure 3B-E). The changes in histone marks coincide with transcriptional changes (Figure 3B-E) and *HOTTIP* binding pattern alteration. Concomitantly with decreased *HOTTIP* binding and active histone marks in *HOTTIP*^-/-^ cells, chromatin accessibility at regulatory loci of all four HOX cluster miRNAs was also reduced (Figure 3G-K). Therefore, *HOTTIP* governs its target genes by regulating the chromatin signature of specific miRNAs in the *MLLr+* AML.

### *HOTTIP* reactivation in *CBS7/9*^*+/−*^ AML cells rescued the miRNA expression coupled with restored chromatin signature

We previously demonstrated that *HOTTIP* coordinates with its upstream regulatory element the CTCF binding site located between *HOXA7* and *HOXA9* (*CBS7/9*) to activate posterior HOXA genes and other hematopoietic oncogenes^11,20^. Reactivation of endogenous *HOTTIP* expression using sgRNA targeted dCAS9-VP160 mediated promoter activation in the *CBS7/9*^*+/−*^ MOLM13 cells efficiently restored posterior HOX gene (*HOXA9-HOXA13*) expression. These findings led us to test whether *HOTTIP* reactivation in *CBS7/9*^*+/−*^ also affects miRNA expression and alters chromatin dynamics at the miRNA loci. Indeed, levels of all HOX cluster miRNAs were significantly repressed in *CBS7/9*^*+/−*^ MOLM13 cells and completely restored in *CBS7/9*^*+/−*^*-HT-VP160* cells with reactivated *HOTTIP* (Figure 4A).

**Figure 4.**
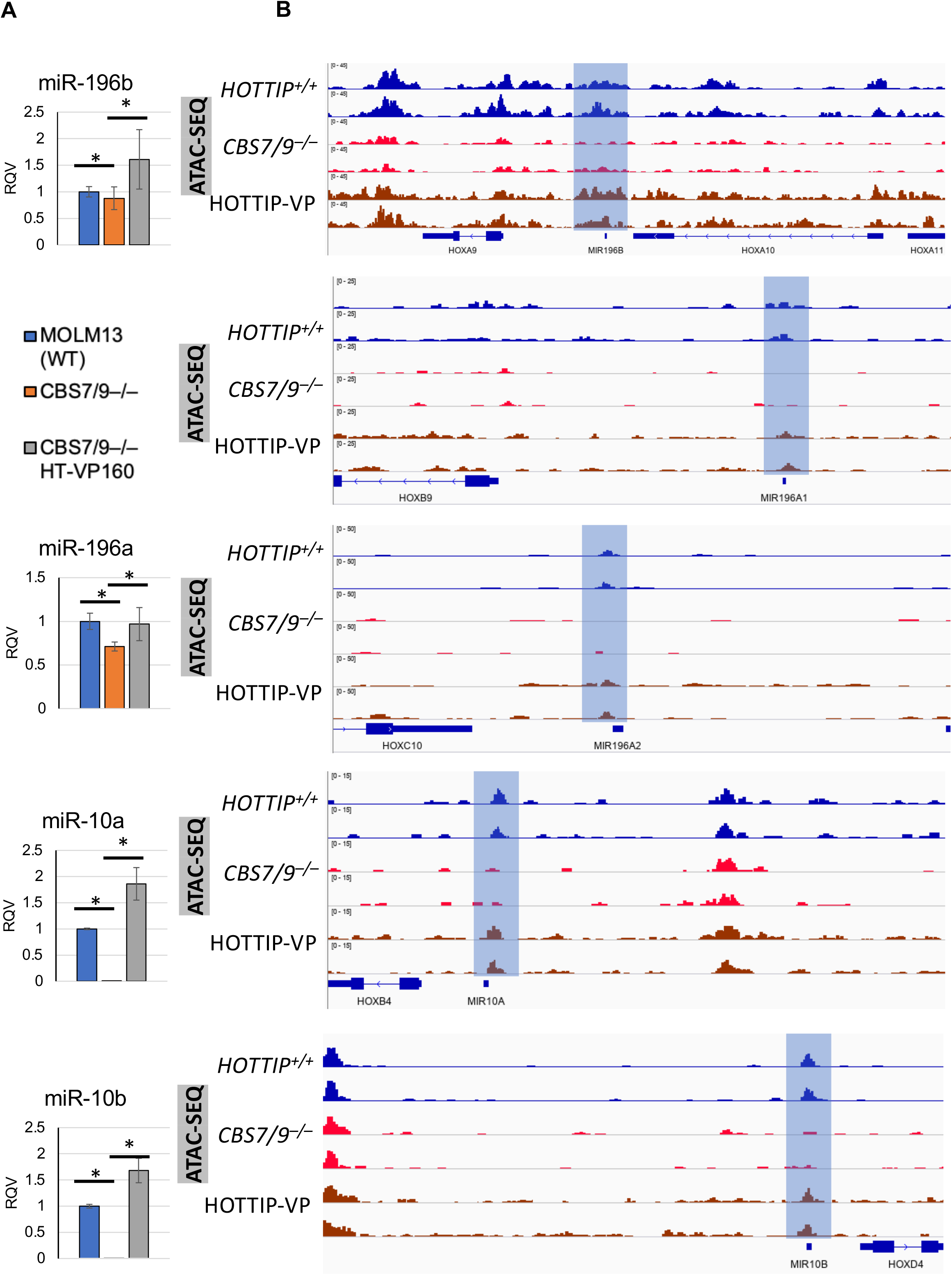
*CBS7/9* boundary regulates HOX cluster miRNAs chromatin neighborhood. (A) qRT-PCR analysis of HOX cluster miRNAs in WT, *CBS7/9*^*+/−*^, and *CBS7/9*^*+/−*^+*HOTTIP*-VP MOLM13 cells. (B) ATAC-seq analysis shows chromatin accessibility on the loci of HOX cluster miRNAs in WT, *CBS7/9*^*+/−*^, and dCas9-VP160-mediated *HOTTIP*-activated MOLM13 cells.

Next, we carried out ATAC-seq data analysis using WT, *CBS7/9*^*+/−*^, and *HOTTIP*-reactivated *CBS7/9*^*+/−*^ MOLM13 cells. Repression of *HOTTIP* in *CBS7/9*^*+/−*^ cell reduced chromatin accessibility at the HOX cluster-associate miRNA loci, while accessibility was largely restored in *HOTTIP*-activated *CBS*^*+/−*^-HT-VP160 MOLM13 (Figure 5B). Hence, *HOTTIP* function is critical for regulating the chromatin accessibility and expression levels of HOX locus miRNAs.

**Figure 5.**
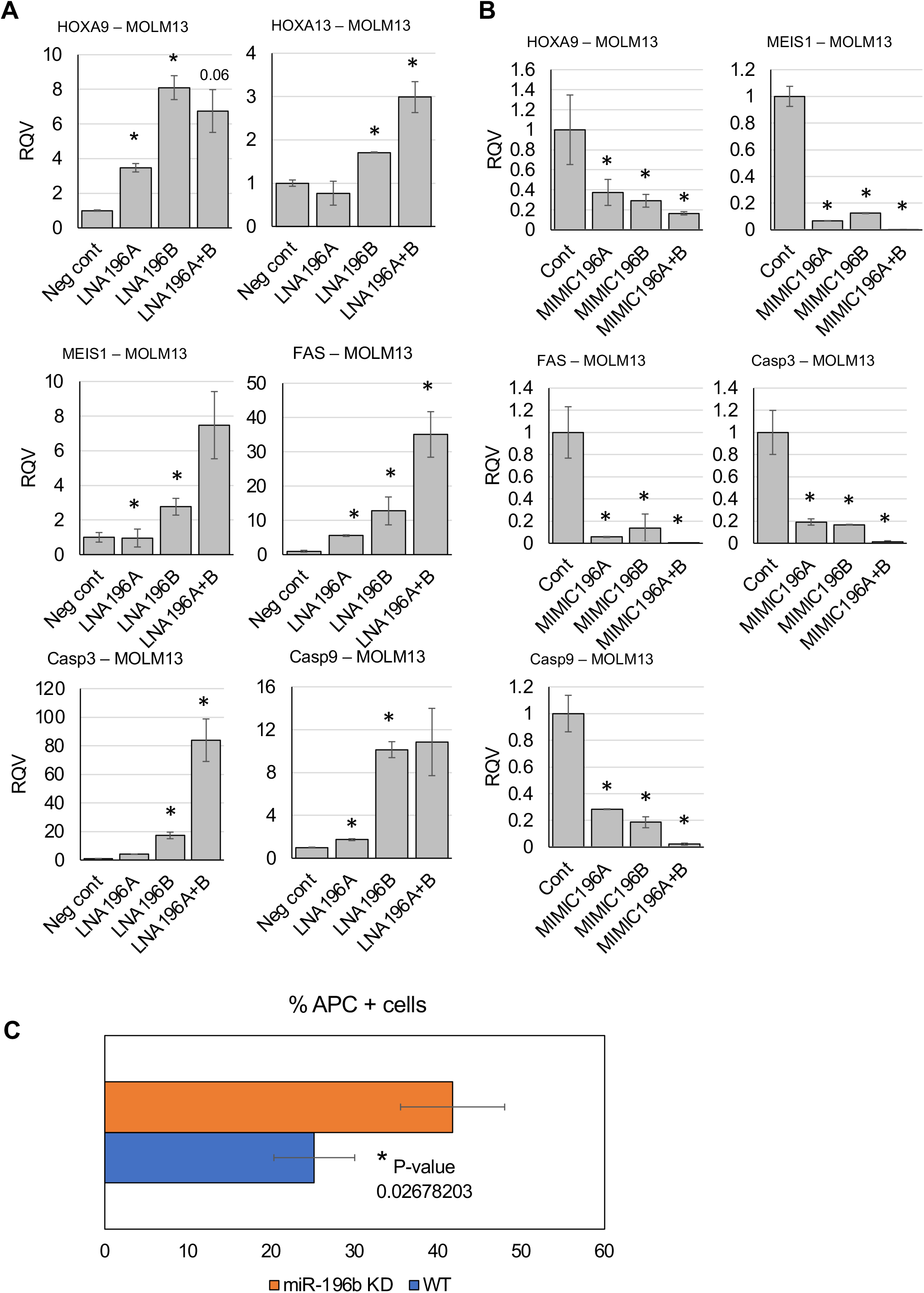
Inhibition of miR-196b induced apoptosis in MOLM13 cells. (A) qRT-PCR analysis of miR-196b targets in MOLM13 cells treated with either negative control or locked nuclei acids (LNA) against miR-196b. (B) qRT-PCR analysis of miR-196b targets in MOLM13 cells treated with either mimics of specific miRNA or control. (C) Bar graph shows FACS (florescence activated cell sorting) evaluated percentage of annexin stained apoptotic population of MOLM13 cells in control and miR-196b knockdown groups.

### miR-196b simultaneously targets oncogenes and tumor suppressors to maintain MLL-AF9 AML

miR-196b represses a subset of targets with tumor suppressor activity *in vivo* and is selectively enriched by cooperation with MLL-AF9 to promote leukemogenesis^18^. Surprisingly, miR-196b also targets *HOXA9 and MEIS1* that play essential oncogenic roles^18^. Because *HOTTIP* positively regulates both miR-196b and its target *HOXA9* and *MEIS1*^11^, we investigated the miR-196b function in leukemogenesis. Expression of miR-196b was strongly suppressed in MOLM13 cells treated with LNA196b compare to NTC (non-targeting control). Although, MOLM13 cells have high levels of *HOXA9* expression, we assessed the regulatory effects of miR-196b on *HOXA9 and MEIS1* in MOLM13 cells. Both *HOXA9* and *MEIS1* were significantly elevated upon miR-196b inhibition (Figure 5a). Reduction of miR-196b also resulted in increased expression of *HOXA13* in MOLM13 cells (Figure 5A). This suggests that repressive function of miR-196b might fine tune expression of *HOTTIP* activated HOX genes in acute myeloid leukemia.

To further understand the pro-oncogenic role of miR-196b, we investigated the expression of tumor suppressor *FAS*, which is a verified target of miR-196b in AML subtypes and colon cancer cells^18,19^. Expression of *FAS* and its downstream genes *Caspase3* (*CASP3*) and *Caspase9* (*CASP9*) was significantly elevated in cells treated with LNA196b compare to NTC (Figure 5A). However, growth competition shows no difference in the proliferation rate of cells treated with LNA196b compared to NTC.

Next, we assessed whether overexpression of miR-196b simultaneously represses the expression of both oncogenes and tumor suppressors. Forced expression of miR-196b significantly represses *HOXA9, MEIS1, FAS, CASP3* and *CASP9* in MOLM13 cells (Figure 5B). To reveal the mechanism underlying the oncogenic role of miR-196b, we analyzed the apoptosis rate in MOLM13 cells treated with either LNA196b or NTC. Repression of miR-196b manifested a higher rate of apoptosis compared to NTC-treated MOLM13 cells (Figure 5C). These results indicate that miR-196b promotes leukemogenesis by down-regulating *FAS,* and its downstream genes, thus suppressing apoptosis in AML.

### *HOTTIP* inhibits *FAS* expression to maintain MLL-AF9 leukemia

We previously reported that enforced expression of *HOTTIP* aberrantly elevated *HOXA9-HOXA13* genes, resulting in impaired hematopoietic stem cell (HSC) function and an increased leukemia stem cell (LSC) population *in vivo*^11^. Thus, we intend to define the mechanisms by which *HOTTIP* might control the LSC pool. Since *HOTTIP* regulates the chromatin signature and expression of miR-196b, which targets both oncogenes and pro-apoptotic genes (e.g., *FAS*) simultaneously, we investigated whether *HOTTIP* also targets FAS in AML. The mRNA level of *FAS* and its downstream pathway genes (e.g., *CASP3*, *CASP8,* and *CASP9*) significantly increased in *HOTTIP*^-/-^ relative to WT MOLM13 cells (Figure 6A). Further, we found that expression of the pro-apoptotic was enhanced in *CBS7/9*^+/−^ cells in which *HOTTIP* expression is significantly repressed (Figure 6B). Intriguingly, when comparing *CBS7/9*^+/−^ cells to WT cells (Figure 6B), mRNA levels of *CASP3* and *CASP9* were significantly elevated in while *CASP8* was unaffected. We next tested whether CRISPR-mediated endogenous gene activation of *HOTTIP* in these cells represses the pro-apoptotic genes. Reactivation of *HOTTIP* in the *CBS7/*9^+/−^ cells strongly repressed *CASP3* and *CASP9* while the *CASP8* level remained unchanged (Figure 6B). Next, we carried out western blot analysis using WT, *HOTTIP*^-/-^, *CBS*^+/−^, and the *HOTTIP*-activated *CBS7/9*^*+/−*^ MOLM13 cells. Cleaved CASP3 protein level was markedly increased in *HOTTIP*^-/-^ and *CBS7/9*^+/−^ compare to WT cells, whereas the Cleaved CASP3 protein level was undetectable upon *HOTTIP* reactivation, which closely resembles WT MOLM13 cells (Figure 6C). Then we examined the elevated level of cleaved CASP3 upon *HOTTIP*^-/-^ induced apoptotic cell death in MOLM13 cells. FACS analysis clearly showed an increase in apoptotic cell death in *HOTTIP*^-/-^ cells compared to WT cells (Figure 6D). Thus, HOTTIP directly controls AML cell survival and apoptosis by regulating the FAS-Caspase axis.

**Figure 6.**
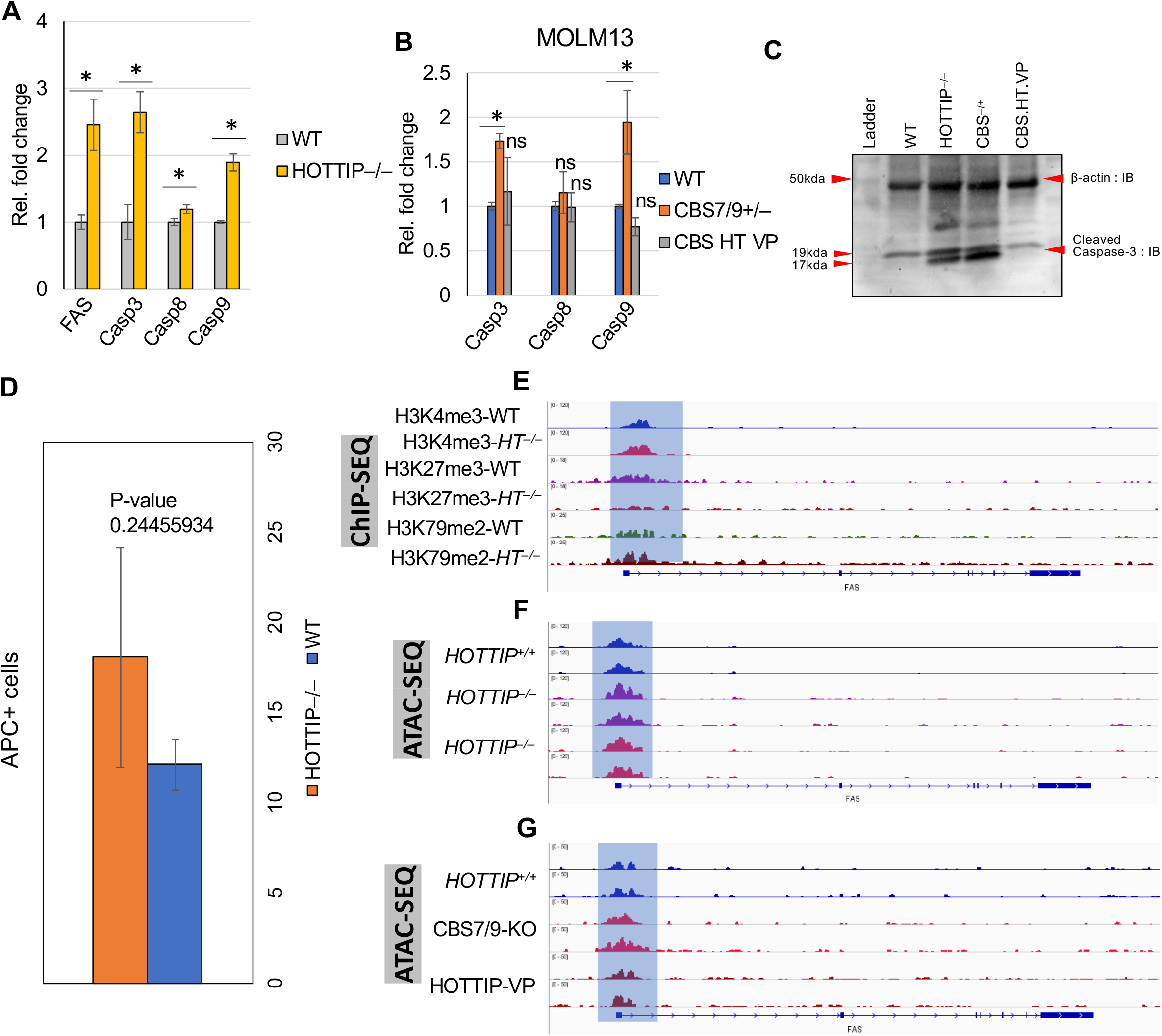
*HOTTIP* targets *FAS,* and *CBS7/9* boundary play role in maintaining the *HOTTIP* targets-chromatin neighborhood. (A) qRT-PCR of the genes associated with apoptosis in WT and *HOTTIP*^-/-^ MOLM13 cells. (B) Expression level of *Caspase* genes in WT, *CBS7/9*^*+/−*^, and dCas9-VP160-mediated *HOTTIP*-activated MOLM13 cells. (C) Western blot shows protein level of ß-actin and Cleaved Caspase-3 in WT, *CBS7/9*^*+/−*^, and dCas9-VP160-mediated *HOTTIP*-activated MOLM13 cells. (D) Bar graph shows percentage of FACS evaluated annexin stained apoptotic population in WT and *HOTTIP*^*-/-*^ MOLM13 cells. (E) ChIP-seq analysis shows histone marks enrichment on the *FAS* promoter in WT and *HOTTIP*^-/-^ MOLM13 cells. (F) ATAC-seq analysis of chromatin accessibility on the *FAS* promoter in WT and *HOTTIP*^-/-^ MOLM13 cells. (G) ATAC-seq analysis of chromatin accessibility on the *FAS* promoter in WT, *CBS7/9*^*+/−*^, and dCas9-VP-160-mediated *HOTTIP*-activated MOLM13 cells.

To evaluate how *HOTTIP* directly regulates *FAS* expression, we investigated the *HOTTIP* binding and chromatin signature at the promoter of *FAS* in WT, *HOTTIP*^-/-^, *CBS7/9^+/−^*, and *CBS7/9*^+/−^HT-VP160 MOLM13 cell lines. Although, enrichment of the *HOTTIP* binding on the *FAS* promoter occurs at relatively low level, *HOTTIP* depletion affects histone modifications associated with active and repressive chromatin. Consistently, relative enrichment of active histone modifications (H3K4me3 and H3K79me2) was increased, whereas repressive histone modification (H3K27me3) was decreased in *HOTTIP*^-/-^ compare to WT cells. Additionally, chromatin accessibility, as determined by ATAC-seq, was increased at the *HOTTIP* binding site on *FAS* in *CBS7/9*^+/−^, whereas reactivation of *HOTTIP* reduced accessibility, resembling WT MOLM13 cells. This suggests that *HOTTIP* maintains a repressive chromatin signature at the promoters of tumor suppressor genes to promote leukemogenesis.

### MiR-196b repression delays leukemogenesis in MOLM13 cell transplantation

miR-196b functions downstream of *HOTTIP*, and together they coordinate to repress *FAS* tumor suppressor in MOLM13 cells. We previously showed that transplantation of *HOTTIP*^-/-^ MOLM13 cells into irradiated NSG mice delayed leukemogenesis compared to WT MOLM13 cells. To further assess the function of miR-196b downstream of *HOTTIP* in leukemogenesis *in vivo*, we knocked down miR-196b in MOLM13-YFP-luciferase cells and then transplanted them into immunodeficient NOD-scid IL2rγnull (NSG) mice for luciferase-based imaging of leukemogenic burden over time and, ultimately, survival analysis. Inhibition of miR-196b inhibited leukemia cell engraftment, whereas transplantation of WT MOLM13-YFP-luc cells showed remarkable expansion *in vivo* (Figure 7A, B, C). We found that depletion of miR196b resulted in elongated survival (19 days, OS), whereas mice receiving WT MOLM13 cells died on day 15 (overall survival, OS) (Figure 7D). The spleen size was dramatically smaller in mice receiving miR-196b KD MOLM13-YFP-luc cells than in mice receiving WT MOLM13-YFP-luc cells (Figure 7E). Fluorescence-activated cell sorting (FACS) analysis showed that CD45+ bone marrow (BM) and spleen cells were drastically reduced in mice receiving miR-196b KD cells compare to WT cells (Figure 7F). Thus, miR-196b functions downstream of *HOTTIP* and coordinates with the lncRNA to repress FAS signaling, and miR-196b inhibition reduces the AML leukemia burden *in vivo* (Figure 7G).

**Figure 7.**
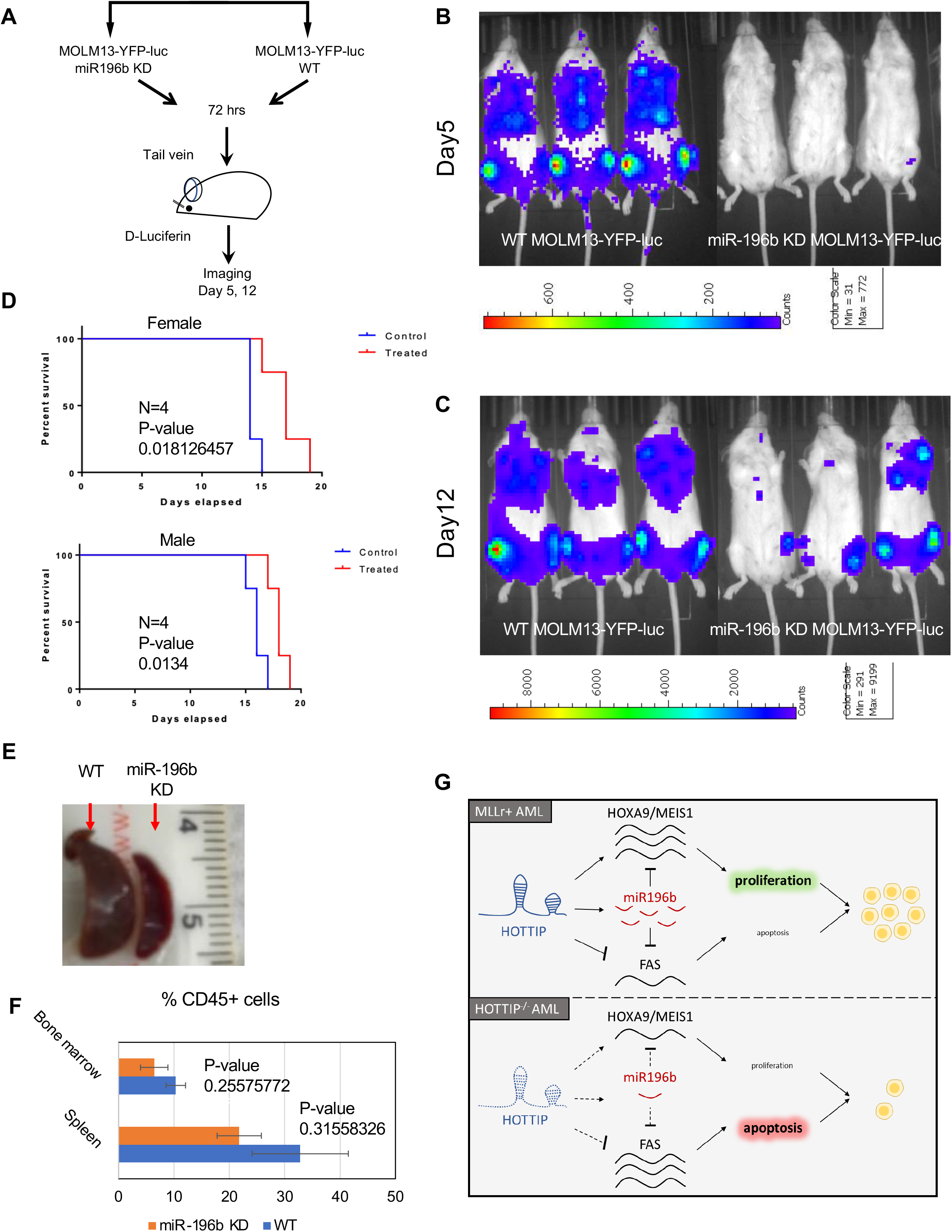
Inhibition of miR-196b inhibits *in vivo* leukemogenesis. (A) Schematic representation of the MOLM13 transplantation in NSG mice and imaging. Luciferase expressing MOLM13 cells either WT or miR196b KD were injected into NSG mice, followed by *in vivo* imaging to assess the knockdown effects of miR-196b in leukemogenesis (B, C) Representative images of leukemogenesis in NSG mice transplanted with MOLM13 cells treated with either LNA196b or negative control. (D) Overall survival (OS) of NSG mice injected with MOLM13 cells. (E) Images of spleens of NSG mice 12 days after transplantion with MOLM13 cells treated with either LNA196B or neg control. (F) Human CD45^+^ cells from bone marrow (BM) and spleen isolated 12 days after transplantation were analyzed by FACS. (G) Model of HOTTIP/miR26b regulation of AML. HOTTIP activation in MLLr+ AML activates expression of *HOXA9*, *MEIS1*, and miR296b and suppresses *FAS*. miR296b, in turn, represses *HOTTIP* target gene to maintain a proliferative state and drive leukemogenesis. In the absence of *HOTTIP*, these pathways are inactive, therefore, *HOXA9* and *MEIS1* levels decrease, and *FAS* levels increase, resulting in leukemic cells undergoing apoptosis.

## Discussion

Long-non-coding RNAs (lncRNAs) and microRNAs (miRNAs) have emerged as biomarkers, drivers, and potential therapeutic targets for a wide array of complex disorders, including leukemia^16^. A growing number of publications demonstrate that miRNAs interact with lncRNAs thereby influencing their target gene expression^21^. The role of lncRNAs in gene silencing is well established; however, less is known about their function in active genomic loci where miRNAs are transcribed^5^. *HOTTIP* (*HOXA* transcript at the distal TIP), a lncRNA, has been known to interact with WDR or WDR/MLL protein complex to epigenetically regulate the 5’ HOXA gene by methylating histone H3K4, which is associated with active transcription. Recent studies from our laboratory combining a variety of molecular biology and bioinformatics techniques have revealed the regulatory networks activated by *HOTTIP* in malignant hematopoiesis and acute myeloid leukemia. Depletion of *HOTTIP* effectively inhibited the posterior HOXA genes (*HOXA9-HOXA13*), and its inhibitory effects gradually diminished on genes towards the anterior end (*HOXA1-HOXA7*)^11,22^. However, whether miRNA genes are regulated by *HOTTIP* in a similar manner to protein coding genes remained a question relevant to hematopoiesis and leukemogenesis. LncRNA can influence miRNA function in many ways: (1) lncRNAs can serve as a source to produce mature miRNAs (2) lncRNAs act as sponge, binding miRNA to prevent them from repressing their target mRNA genes and (3) lncRNA alter miRNA gene transcription by binding on their promoter^16^. To profile *HOTTIP* regulated miRNA in MOLM13 cells, we carried out small RNA sequencing that revealed that loss of *HOTTIP* significantly affects expression of several miRNAs when comparing WT and *HOTTIP*^-/-^ MOLM13 AML cells. Intriguingly, several ncRNAs, including lncRNAs and miRNAs, are transcribed from the intergenic regions of the HOX clusters. However, it remains unknown how *HOTTIP* may mediate ncRNA transcription, particularly of miRNAs, and how the *HOTTIP-*miRNA axis governs gene expression in malignant hematopoiesis.

*HOXA9* expression is positively regulated by mixed lineage leukemia (MLL) methyltransferase, which trimethylates histone 3 lysine 4 (H3K4me3) at the *HOXA9* promoter^23^. This mechanism is directly antagonized by the sequential activity of polycomb repressive complexes PRC1 and PRC2 that trimethylated histone 3 lysine 27 (H3K27me3). Moreover, topologically associating domains (TAD) of the HOX loci within the nucleus also have an important role in coordinating expression. Long non-coding RNA *HOTTIP* interacts with the WDR5-MLL complex and localizes the complex to the 5’*HOXA* locus. To determine whether *HOTTIP* provides a basis for transcriptional activation and three-dimensional (3D) chromatin organization in posterior HOX gene loci, we screened all CTCF sites and lncRNAs important for *HOXA9* expression within all four HOX gene loci in *MLL-AF9* rearranged MOLM13 AML cells using a CRISPR/CAS9 lentivirus screening library^11,24^. The *HOTTIP* lncRNA acts downstream of the *CBS7/9* boundary to regulate expression of genes located in *cis* and *trans*, including *HOXA13-HOXA9, MEIS1,* and *RUNX1*, which are important for hematopoiesis, and leukemia^11^. Furthermore, ChIRP-seq analysis revealed that *HOTTIP* binds to its target genes^11^ and a cohort of miRNAs. This data raises the question of whether *HOTTIP* also controls the expression of miRNAs that are involved in the management of malignant hematopoiesis.

Our analysis demonstrated that inhibition of *HOTTIP* in MOLM13 cells significantly inhibited several miRNAs, including HOX cluster miRNA, miR-196b, which targets both oncogenic *HOXA9* and the *FAS* tumor suppressor. Expression profiling of primary AML patient samples showed strong correlation among miRNA, *HOTTIP*, and their target HOX genes. Furthermore, ATAC-seq (assay for transposase-accessible chromatin using sequencing) data analysis defined chromatin signatures at differentially expressed miRNAs bound by *HOTTIP* in *cis* and *trans*, including HOX clusters miR-196b. *HOTTIP* binding on the promoter region of miRNAs that are differential expressed in *HOTTIP*^-/-^ MOLM13 cells suggests that *HOTTIP* directly controls transcriptional regulation of miRNAs.

Dynamics of chromosomal structure play important roles in gene control. A number of proteins modulate chromatin dynamics by contributing to structural interactions between gene promoters and their enhancer elements. Enhancer/promoter communications for specific transcription programs are enabled by topological associated domains (TADs), which are basically structural and functional chromosomal units. Often inappropriate promoter/enhancer interactions result from altered TADs, which lead to aberrant transcription of oncogenes or tumor suppressor genes. Binding of transcription factor CTCF in chromatin boundaries plays an important role in defining TADs and chromatin signature within TADs. We previously reported that CTCF binding located in between of *HOXA7* and *HOXA9* defines posterior HOXA locus TADs and chromatin signature within the TADs. Deletions of *CBS7/9* impaired chromatin structure and altered posterior HOXA gene expression due to lacking function of *HOTTIP* lncRNA^11^. By virtue of *CBS7/9’s* role in regulating posterior HOXA genes and lncRNA *HOTTIP* expression, we show that HOX cluster miRNAs are altered in *CBS7/9*^*-/-*^ and that *HOTTIP* over-expression restored *CBS7/9*-mediated HOX cluster miRNA expression and chromatin signature. Apart from the cis coding miR-196b, *HOTTIP* lncRNA also bound and regulated *trans* HOX and non-HOX cluster miRNAs.

To decipher the mechanisms by which *HOTTIP* exerts miRNAs to control target genes, we performed bioinformatics analysis that revealed a large number of differentially expressed genes in *HOTTIP*^-/-^ MOLM13 cells, which are co-regulated by *HOTTIP* and miR-196b. Although *HOTTIP* positively regulates expression of their co-expressed oncogenes, several of them are negatively regulated by miR-196b. Thus, it seems that negative regulation of the co-expressed 5’ HOXA genes in MOLM13 cells might be fine-tuned by miR-196b in normal hematopoiesis^18^. Notably, single miRNA (or groups of miRNAs) target multiple genes including oncogenes and tumor suppressors simultaneously or sequentially. Partial repression of *HOXA9* and *MEIS1* by miR-196b in the human *MLL*-rearranged leukemia may not potent enough to affect their function to induce and maintain leukemia. The tumor suppressor targets (e.g., *FAS*) of miR-196b could significantly inhibit cell transformation and leukemogenesis. Indeed, miR-196b inhibition induced *FAS* expression and *Cleaved Caspase-3*. As a result, the apoptotic cell death increased upon miR-196b knockdown in MOLM13 cells. Similarly, induced expression of *FAS* and *Cas3* in *HOTTIP*^-/-^ and *CBS7/9*^*-/-*^ is associated with increased cell death. *HOTTIP* modulates epigenetic marks on the *FAS* promoter and thereby controls chromatin accessibility and gene expression. However, *HOTTIP* represses *FAS* expression perhaps through activation of miR-196b that directly targets *FAS*. miR-196b-deficient MOLM13 cell transplantation in NSG mice delayed leukemogenesis. All mice transplanted with WT MOLM13 cells died within 15 days, while mice receiving miR-196b-depleted cells survived longer and exhibited fewer CD45+ cells. Transgenic overexpression of *HOTTIP* lncRNA in mice affected HSC function and increased leukemia stem cell (LSC) pool, inducing leukemia-like disease^11^. Thus, *HOTTIP* and miR-196b deletion reduces the AML leukemic burden *in vivo*, and both coordinate to regulate *FAS* expression at the transcriptional and post-transcriptional level to promote leukemogenesis.

Taken together, we report a mechanism mediated by *HOTTIP* to regulate miR-196b expression in AML. Our study revealed a regulatory model in which *HOTTIP*-miR-196b axis repress expression of tumor suppressor *FAS* that circumvent the negative effects of *HOXA9* repression by miR-196b in AML. The aberrant upregulation of both *HOTTIP* and miR-196b by MLL fusion results in the persistent repression of its tumor-suppressor targets (e.g., *FAS*) along with dual control (transactivation and inhibition) of their oncogenic co-targets (*HOXA9/MEIS1*). This inhibits differentiation, disrupts cell homeostasis, and promotes cell proliferation via inhibiting apoptosis, consequently maintaining leukemia stem cell pools. Apart from miR-196b, *HOTTIP* also bound and regulated a subset of non-HOX cluster miRNAs while a subset of miRNAs was upregulated in *HOTTIP*-inhibited MOLM13 cells. Future studies should aim to investigate the mechanism by which *HOTTIP* modulates expression of candidate miRNA in hematopoiesis vs leukemia.

## Supplementary Figures

**Supplementary Figure 1.**
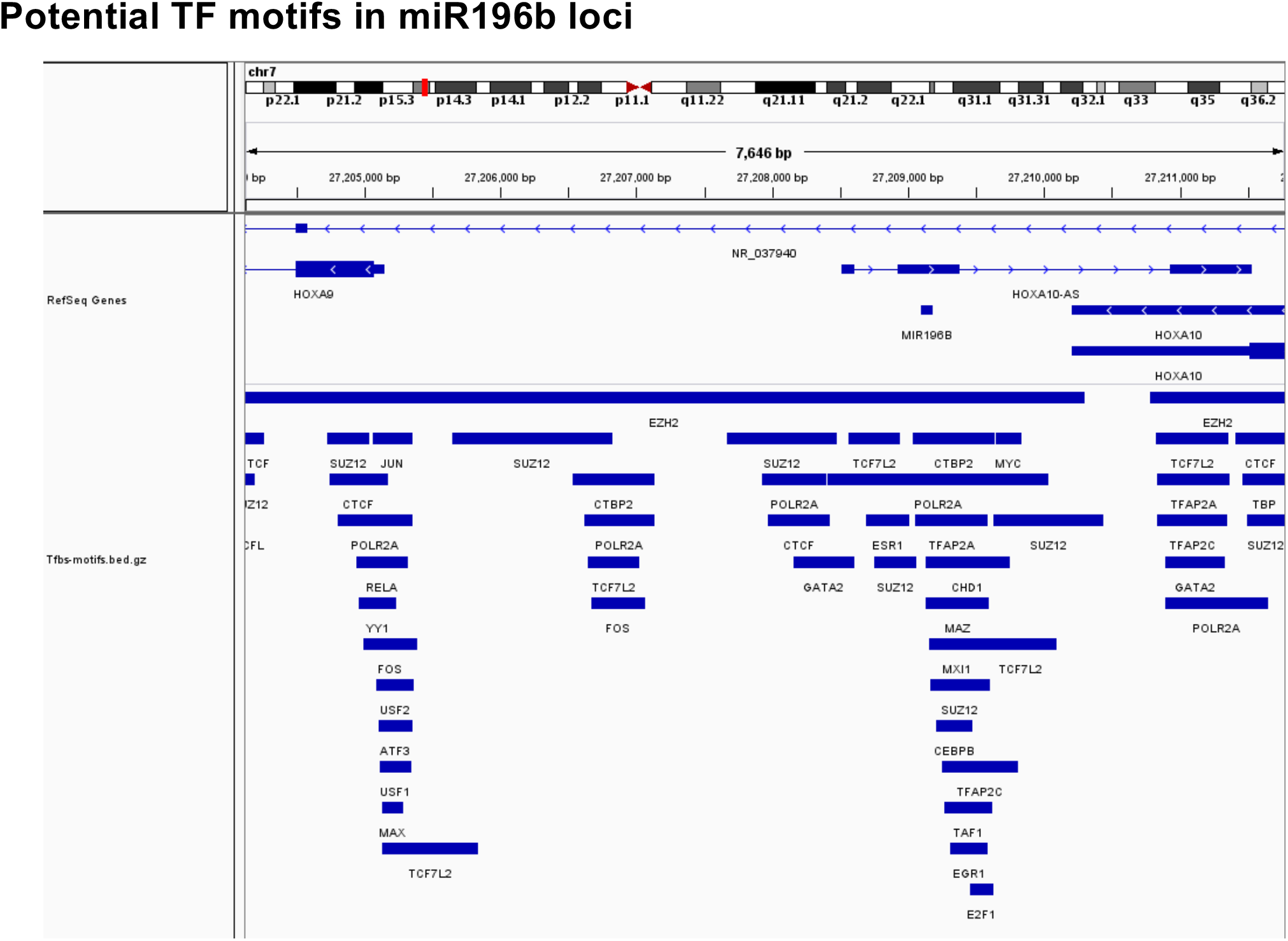
UCSC gene track shows transcription factor binding motifs on the miR-196b genomic locus and posterior HOXA genes.

**Supplementary Figure 2.**
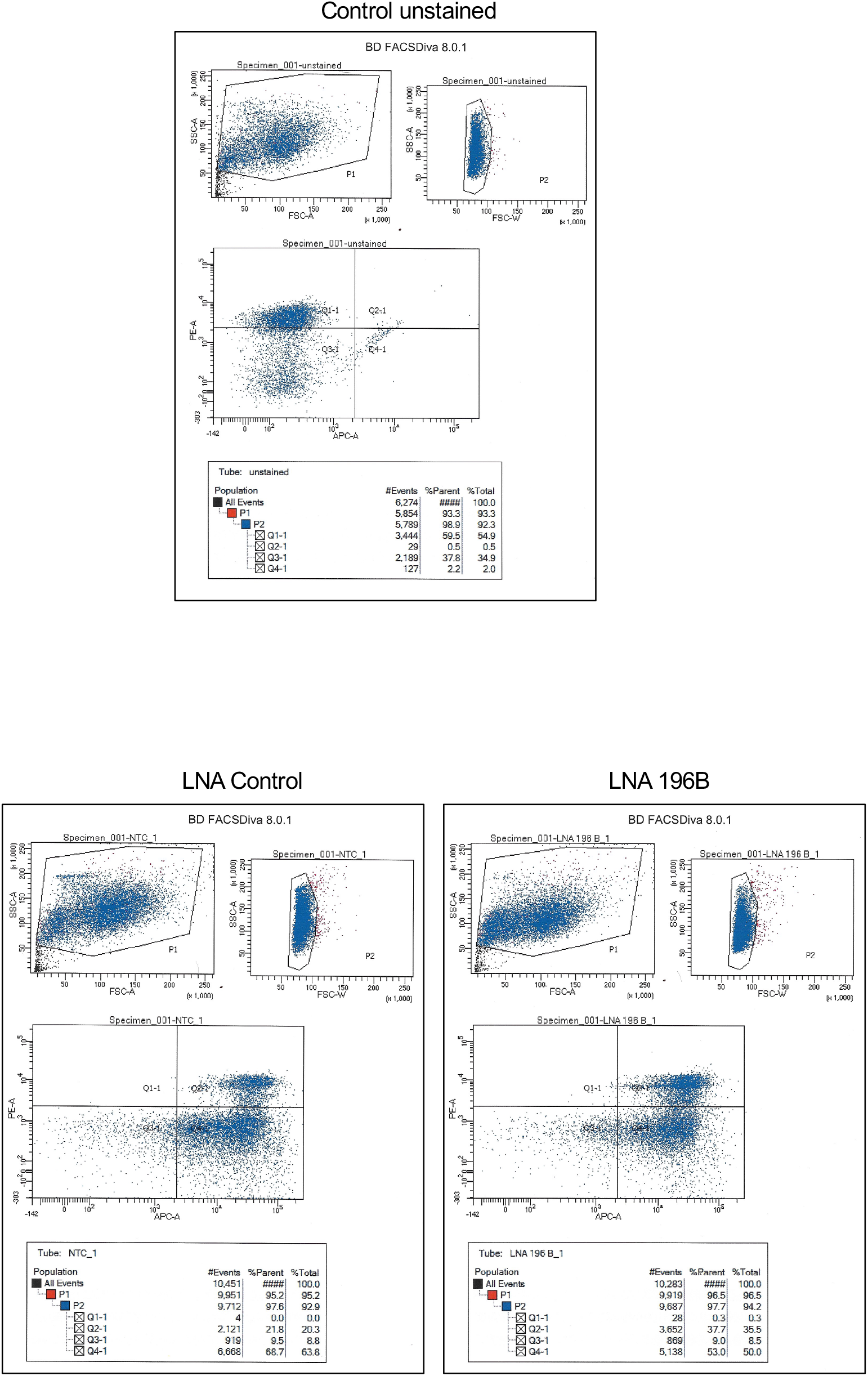
FACS analysis of annexin stained cells upon knockdown of miR-196b, and control.

**Supplementary Figure 2.**
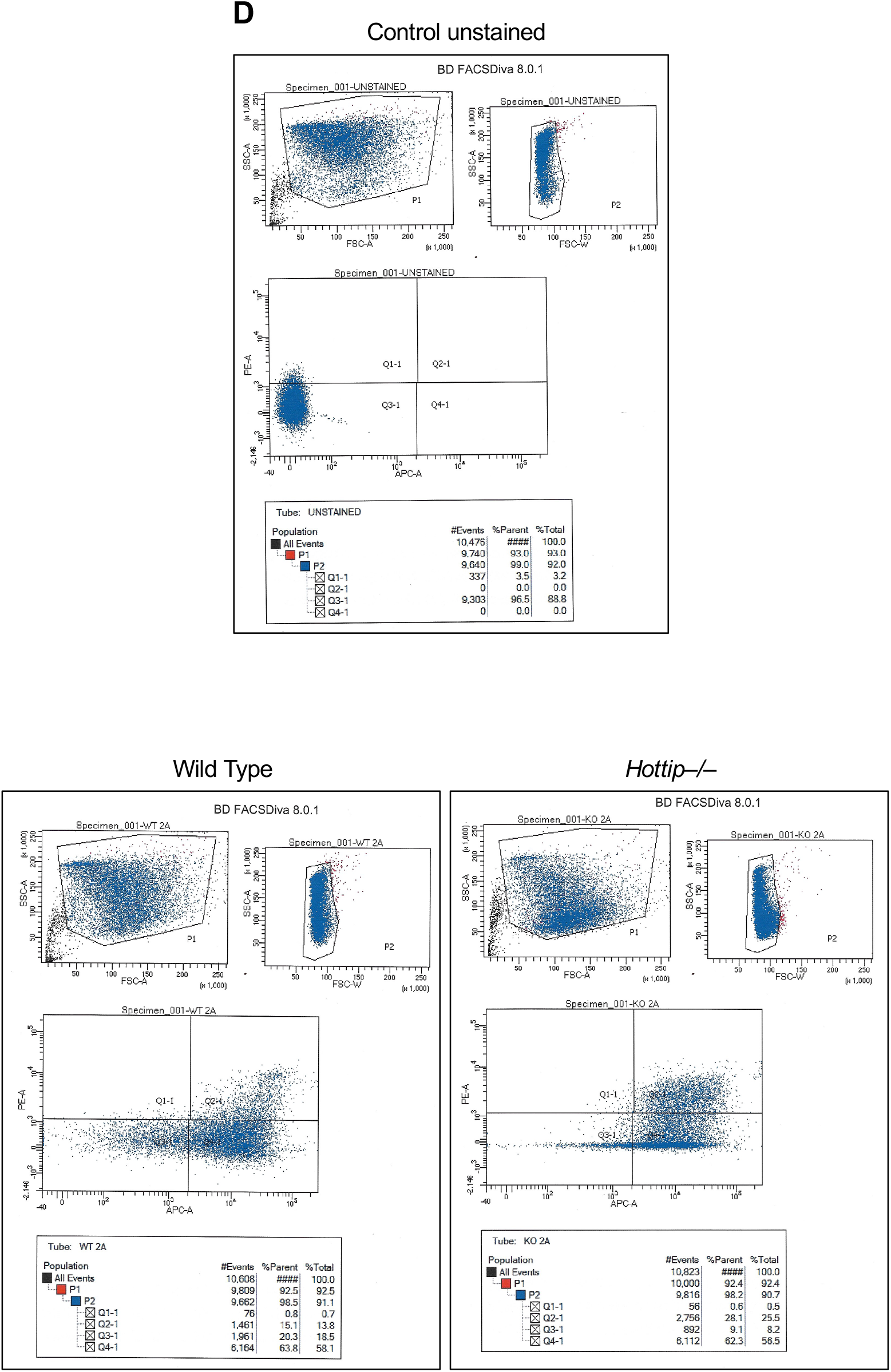
FACS analysis of annexin stained *HOTTIP*^-/-^ and wild type MOLM13 cells.

**Supplementary Figure 2.**
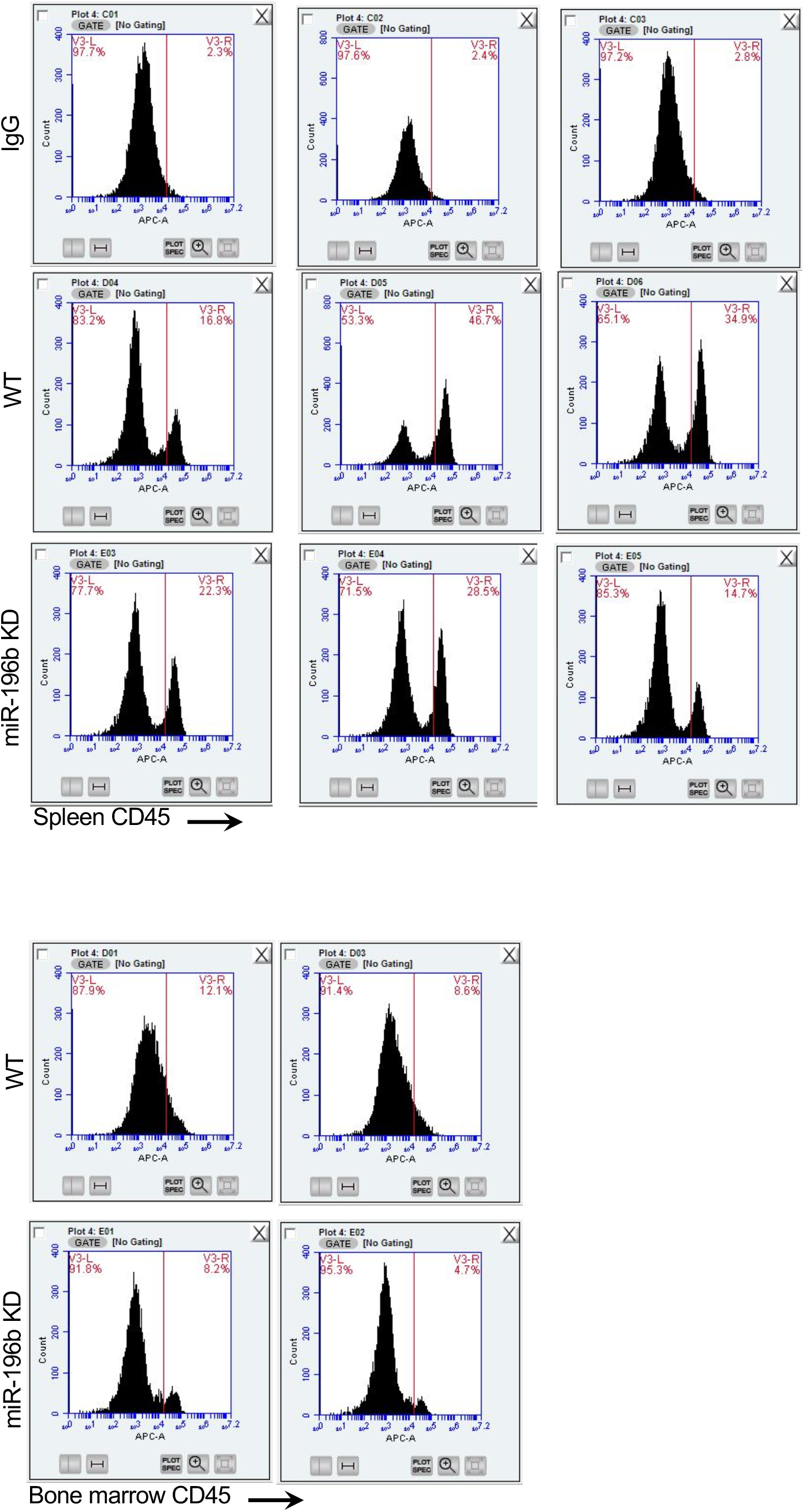
FACS analysis of CD45 stained bone marrow and spleen cells, harvested from NSG mice transplanted with either WT or miR-196b knocked down miR-196b MOLM13 cells.

## Material and Methods

### Tissue and Cell culture

MOLM13 cells were cultured and maintained in RPMI medium supplemented with 10% FBS and 1% penicillin and streptomycin solution. All medium and supplements were purchased from Thermo Fisher Scientific.

### SsecCRISPR-Cas9 Mediated *HOTTIP* lncRNA Knock-Out and Lentivirus Production

The detailed protocol of generating *HOTTIP* knockout (KO) MOLM13 leukemia cells were described previously in Luo et. al. 2019. In brief, CRISPR-RNA (crRNA) and tracrRNA were mixed and annealed at 95^0^C for 5 min and the cooled down to room temperature. Subsequently, crRNA: tracrRNA duplex and S.p. Cas9 Nuclease components were combined together and then mixed with 500,000 AML cells for electroporation with Neon^®^ System. The DNA was extracted after 24 hrs. or 96hrs. from 100ul of transfected cells using Qiagen Quick Extract kit and processed for Sanger sequencing for verification of mutation. The targeted deletion of *HOTTIP: HOTTIP*^*-/-*^ −#1 targeted region is Chr7: 27241953-27241985; *HOTTIP*^*-/-*^ −#2 targeted region is Chr7: 27240098-27240123.

### dCas9-Mediated Overexpression of *HOTTIP* in AML Cells

*HOTTIP* promoter targeting guide RNA was designed using the Zhang laboratory web tool (http://crispr.mit.edu, and cloned into lentiviral vector the pLKO5.sgRNA.EFS.tRFP vector (Addgene #57824). The gRNA plasmid encoding puromycin was co-transfected with a plasmid encoding dCas9-VP160 (pAC94-pmax-dCas9VP160-2A-puro, addgene plasmid number #48226) in MOLM13 and OCI-AML3 cells. Transfected MOLM13 cells were selected with 2ug/ml puromycin for 48 hrs. post-transfection, and then FACS sorted for RFP^+^ cells. RNA was extracted from RFP^+^ cells, and gene expression was analyzed by qRT-PCR using specific primer sets (Luo et al. 2019).

### MiRNA Knockdown and Over Expression

Locked Nucleic Acids (LNA) and miRNA mimics and scrambled LNA/mimic (negative control) was purchased from Qiagen and MOLM13 cells were transfected with either LONZA nucleofector devise (program X-001) or Lipofectamine 3000 (Invitrogen). Cells were harvested post 72 hrs. of transfection for RNA extraction and gene expression analysis.

### Western Blot Analysis

Whole cell lysate extract (total protein) was prepared using RIPA buffer and quantified using Bradford method. Total protein lysates were fractionated on 4%-20% on tris-glycine polyacrylamide gradient gel and transferred onto PVDF membrane. The bot was exposed to specific antibodies to detect the endogenous protein level using chemiluminescence method and BioRad imaging system: anti-ß-actin (A2066-100U); Sigma at 1: 5000, anti-Cleaved Caspase-3 (9661); Cell Signaling at 1:1000 dilution.

### Apoptosis Analysis

MOLM13 cells transfected with either LNA against specific miRNA or scrambled non-targeting control were seeded and cultured in 6-well dishes. Cells were harvested 72hrs post transfection and washed with PBS. Apoptotic cells were detected by Accuri C6 fluorescence-activated cell sorting (FACS) using Annexin V-APC Apoptosis Detection Kit (BD #), according to the manufacturers protocol.

### RNA Extraction and Real-Time qPCR

Norgen Biotek RNA purification kit was used to isolate total RNA as per manufacturer’s instructions. cDNA (reverse transcription - RNA to cDNA) was made using High Capacity RNA to cDNA kit (Life Technologies, Grand Island, NY). miRNA and mRNA qPCR were performed using TaqMan (Life Technologies) and Sybr Green assay respectively with either Taqman Universal PCR Master Mix (miRNA qPCR) or BioRad SsoAdvanced Universal Sybr Green Supermix according to the manufacturer’s protocol on a Bio-Rad CFX96 Touch Real Time PCR Detection System (Bio-Rad Laboratories, Richmond, CA). PCR reactions were performed in triplicate using either U6 (miRNA qPCR), or GAPDH (human messenger RNA qPCR) as the normalizer.

### Small RNA library preparation and sequencing analysis

The small RNA-seq of the MOLM13 WT and *HOTTIP*^-/-^ cells was conducted at Genome science Facility, Pennsylvania State University College of Medicine. Small RNA-seq. libraries were prepared using BioO from Perkin Elmer and Qiagen kits. Small RNAs were sequenced using a TruSeq Small RNA Sequencing Kit (Illumina, San Diego, CA, USA) according to manufacturer instructions. The quality of libraries was assesses based on size distribution and concentration using 2100 Bioanalyzer with DNA 1000 chip (Agilent Technologies. All samples were sequenced 25M reads on an Illumina NovaSeq 6000 Sequencer using the 2×50bp paired-end platform.

All of raw binary base call files from the sequencer were transformed into FASTQ format and de-multiplexing using Illumina bcl2fastq2 Conversion Software v2.20 (https://support.illumina.com/downloads/bcl2fastq-conversion-software-v2-20.html).

Quality of the sequenced reads was evaluated using FastQC developed by Babraham Bioinformatics (https://www.bioinformatics.babraham.ac.uk/projects/fastqc/). Then these fastq files were performed by adapter trimming using the FASTQ Toolkit App version 1.0 of Illumna BaseSpace (http://basespace.illumina.com/apps/) and sequence alignment with GRCh37 human genome assembly database using OASIS2.0 (https://oasis.dzne.de/index.php).

Differential expression analysis was performed using the DESeq2 algorithm ^25^ and the expression was normalized using de Variance Stabilizing Transformation from the DESeq2 algorithm in R. The differential expression of miRs with adjusted *p* values < 0.05, and fold change (FC)≥2 representing positive log2FC (>1.0) and negative log2FC (<−1.0) were considered to be significantly different.

The potential targets of miRs were derived from the miRTarBase ^26^ in human genome (http://mirtarbase.cuhk.edu.cn/php/index.php). Raw sequencing data and miRNA quantification tables can be accessed through GEO record #.

### Chromatin Immunoprecipitation (ChIP) Assay

ChIP were performed as described in Luo et. al., 2019. Briefly, chromatin prepared from MOLM13 cells were immunoprecipitated with antibodies against MLL1, H3K4me3, H3K9me2, H3K27me3, and H3K79me2, separately. The MLL1, H3K4me3, H3K79me2 and H3K27me3, H3K79me2 ChIP-DNA libraries were prepared using illumina’s TruSeq ChIP Sample Preparation Kit according to the manufacturers’ instructions (Cat # IP-202-1012). Agilent TapeStation was used to check the quality of the libraries as per manufacturer’s instruction. Final libraries were submitted to paired-end sequencing of 100 bp length on an Illumina HiSeq 3000.

### Chromatin Isolation by RNA Immunoprecipitation (CHIRP) Assay

The detailed methods of CHIRP assay were described in Luo et. al. 2019, briefly sonicated chromatin materials diluted using hybridization buffer and hybridized with 100pmol of biotinylated DNA probes targeting *HOTTIP* or *LacZ* containing 100 mL of Streptavidin-magnetic C1 beads (Invitrogen). RNA and DNA hybrids and RNA binding proteins were subjected to analysis by qRT-PCR and western blot with respective antibodies respectively. Illumina’s TruSeq ChIP Sample Preparation Kit was used according to manufacturer’s instructions for preparation of CHIRP libraries and submitted to paired-end sequencing of 100 bp length on an Illumina HiSeq 2500. All sequencing genomics datasets were deposited in the NCBI GEO under accession number (GSE114981).

### ChIP-seq and ChIRP-seq Data Analysis

ChIP-seq and ChIRP-seq data processing and analysis was described in Luo et. al. 2019. Briefly, all sequencing tracks were viewed using the Integrated Genomic Viewer (Robinson et al., 2011). Peaks annotation was carried out with the command ‘‘annotatePeaks.pl’’ from HOMER package (Heinz et al., 2010). For ChIRP-seq motif analysis, the *de novo* motif analysis was performed by the ‘‘findmotifsgenome.pl’’ from the HOMER motif discovery algorithm (Heinz et al., 2010). The genes and pathways regulated by the *HOTTIP* bound promoters or intergenic regions were analyzed and annotated by the Gene Ontology analysis with the Database for Annotation, Visualization and Integrated Discovery (DAVID) tool (https://david.ncifcrf.gov/, Version 6.8) (Huang da et al., 2009a; Huang da et al., 2009b). Each GO term with a p value more than 1 × 10^-3 is used for cutoff (threshold: 10^−3). All genomics datasets were deposited in the NCBI GEO under accession number (GSE114981).

### Assay for Transposase-Accessible Chromatin Using Sequencing (ATAC-seq)

ATAC-seq was previously described in Luo et. al. 2019 using the Nextera DNA library preparation kit. Libraries were quantified using qPCR (Kapa Library Quantification Kit for Illumina, Roche), and purified with AMPure beads (Beckman Coulter). Quality of the DNA library was examined by Agilent bio-analyzer 2100 prior to sequencing in duplicates with 2×100 bp paired-end reads on an Illumina NextSeq 500.

### ATAC-seq Analysis

Detailed method of ATAC-seq analysis was previously described in Luo et. al. 2019. Briefly, all sequencing tracks were viewed using the Integrated Genomic Viewer (IGV/2.4.19) (Robinson et al., 2011). Peaks annotation was carried out with the command ‘‘annotatePeaks.pl’’ from HOMER package (version 4.8) (Heinz et al., 2010) and GREAT (McLean et al., 2010). DEseq2 (Benjamini-Hochberg adjusted p< 0.05; FoldChangeR2) were also performed to find the differential binding sites between two peak files, including control and treatment groups with C+G normalized and ‘‘reads in peaks’’ normalized data (Ross-Innes et al., 2012). The *de novo* motif analysis was performed by the ‘‘findmotifsgenome.pl’’ from the HOMER package (Heinz et al., 2010). For each genomic feature (peaks or chromVAR annotation), we calculated the chromatin accessibility median deviation z-score (for chromVAR features) or fragment counts (for peaks) in control and treatment groups with chromVAR package in R language (Rubin et al., 2019; Schep et al., 2017). Pearson’s correlation coefficient and Pearson’s c2-test were carried out to calculate overall similarity between the replicates of ATAC-seq global open chromatin signatures. All genomics datasets were deposited in the NCBI GEO under accession number (GSE114981).

### Xenotransplantation of Human Leukemic Cells and Patient-Derived Xenografts (PDX)

MOLM13 cells - WT or miR-196b KD were injected into the tail vein of the NSG mice (2-3 months old). Cells were resuspended into DPBS and injected at 5 × 10^5^ cells (in 150 to 200ul DPBS) per mouse. Mice were monitored daily for symptoms of disease (ruffled coat, hunched back, weakness and reduced motility) to determine the time of killing for injected mice with sign of distress. Experimentally used NSG mice were humanely killed at the time of moribund. Blood was collected into microtubes for automated process with K_2_EDTA. bone (tibias, femurs and pelvis) and spleen were dissected. BM cells were isolated by flushing the bones. Spleens were mashed through a 70-mm mesh filter and made into single cell suspensions. Human CD45 (BD #) chimerism in these hematopoietic tissues was analyzed by Accuri C6 flow cytometry.

## Acknowledgements

This work was supported by the grants from National Institutes of Health (S.H., R01DK110108, R01CA204044, HL141950)

## Author Contributions

A.P.S., H.L., and S.H. conceptualized the study and formal data analysis. A.P.S. designed and performed the experiments. H.L. and M.E. contributed bioinformatics analysis. M.M. helped with qRT-PCR. A.P.S. and A.S. performed mouse transplantation studies. Writing-original draft, A.P.S., Review and editing, A.P.S., H.L., M.E., S.H., Visualization, A.P.S., H.L., M.E., S.H., Project supervision, A.P.S. and S.H...

## Notes

### Competing Interest Statement

The authors have declared no competing interest.

